# The selfing syndrome overshadows other differences when comparing fitness across *Capsella* species

**DOI:** 10.1101/2020.11.26.398016

**Authors:** Marion Orsucci, Theofilos Vanikiotis, Maria Guerrina, Tianlin Duan, Sylvain Glémin, Martin Lascoux

## Abstract

Self-fertilization has recurrently evolved from outcrossing. Self-fertilization provides an advantage in the short-term as individuals do not require a mate to reproduce, but self-fertilization is also associated with both decreased genetic diversity and accumulation of weakly deleterious mutations, which could, however, be alleviated in polyploid selfers. If pollinators are not limited, individual fitness is thus expected to be higher in outcrossers than in selfers. We measured several life history traits in four *Capsella* species under two different treatments (disturbed and undisturbed) to assess the effects of mating system and ploidy level on reproductive, vegetative and phenological traits. The experiment was carried out outdoor in Northwest Greece, within the range of the obligate outcrossing species, *C. grandiflora*, so it could be naturally pollinated and its fitness directly compared to that of its self-fertilizing relatives. Disturbance of the environment did not affect the phenotype in any of the four species. However, for most traits the obligate outcrossing species performed better than all selfing ones. In contrast, polyploidy did not seem to confer an advantage in terms of survival or reproduction compared to diploidy. Finally, plants from Asia and northern Europe had lower performances than accessions from southern Europe and the Middle-East.

## INTRODUCTION

Mating system shifts and polyploidization events are two major evolutionary transitions in plant evolution that can eventually lead to speciation by promoting both reproductive isolation and ecological divergence (Otto & Whitton, 2000; Wright *et al*., 2013).

Mating system shifts from outcrossing to self-fertilization occurred repeatedly during plant evolution, most likely since it provides reproductive assurance and because of the transmission advantage of selfing mutations (Stebbins, 1974; Barrett, 2002). Prezygotic barriers due to the isolation of the mating system and associated floral differences (Kiang & Hamrick, 1978; Diaz & Macnair, 1999; Sicard & Lenhard, 2011; Wright *et al*., 2013) could reduce gene flow and lead to speciation. Outcrossing and selfing sister species may also diverge ecologically: selfing plants do not depend any longer on pollinators and thereby could be better at colonizing new environments (‘Baker Law’; Pannell, 2015), selfing is often associated with invasiveness (Van Kleunen *et al*., 2008) and selfing species tend to have larger geographic ranges than their outcrossing relatives (Van Kleunen & Johnson, 2007; Randle et al. 2009, Grossenbacher *et al*., 2015). In contrast outcrossers could be better competitors (Munoz *et al*., 2016) and would not suffer from the negative effects of inbreeding (lack of genetic diversity, accumulation of deleterious mutations; Stebbins, 1957; Wright *et al*., 2008; Glémin & Galtier, 2012; Igic & Busch, 2013). Consequently, in a given environment where pollinators are not limited, outcrossing species should have a higher fitness than their self-fertilizing relatives, and this difference should be enhanced by competing conditions.

Polyploid self-fertilizing species make this simple model somewhat more complex. Polyploidization, the increase in genome size caused by the inheritance of an additional set of chromosomes, may also lead to instant speciation. The resulting hybrid will have a high degree of post-zygotic reproductive isolation from its progenitors: backcrossing to either parent will produce nonviable or sterile offspring (Grant, 1981; Ramsey & Schemske, 1998). Two types of polyploid species can be distinguished: (i) autopolyploids originating from whole genome duplication and (ii) allopolyploids that result from the fusion of the genomes of two different diploid species into a unique individual. Polyploidy is often associated with selfing (Barringer, 2007; Robertson *et al*., 2011) and could favor the evolution of selfing by breaking down self-incompatibility systems (Mable, 2004) and reducing the impact of inbreeding depression (Ronfort, 1997). Polyploidization could mitigate the deleterious effects of selfing due to a transient advantage in masking deleterious mutations (Otto & Whitton, 2000). Indeed, the evolutionary success of polyploids has often been ascribed to their greater fitness relative to their diploid relatives (an idea dating back to Shull 1929) and the acquisition of new properties (e.g. salt resistance) although fitness advantage over their diploid relatives has seldom been thoroughly investigated (Blanc & Wolfe, 2004).

In the case of both transitions, shift in mating system and polyploidization, research has focused on the new traits that were acquired. For instance, a large amount of work has been devoted to the genetic basis of changes in floral morphology during the transition from outcrossing to selfing (Sicard & Lenhard, 2011) and on new properties (e.g. salt tolerance, resistance to biotic and abiotic stresses; Levin, 1983; Chao *et al*., 2013) conferred by polyploidy. Less attention has been paid to the traits that may have been retained from the ancestors. However, there are good reasons to believe that both polyploids and selfers might differ less than expected from their ancestors, and in particular retain some of their main ecological properties. In allopolyploids, Gottlieb (2003) coined the term “parental legacy” to describe the fact that properties of the two inherited genomes were preserved to a variable extent. In *Capsella bursa-pastoris* we recently showed that parental legacy in term of polymorphism and gene expression level was indeed important (Kryvokhyzha *et al*., 2019a,b). It is still unclear, however, how much the species are affected by the ploidy levels and/or mating systems in term of ecological strategy and fitness.

The *Capsella* genus (Brassicaceae) has some distinct advantages to address such questions. It is a small and well delineated genus with no documented hybridization with other species (Hurka & Neuffer, 1997; Hurka *et al*., 2012). Yet, the *Capsella* genus combines species with different ploidy levels and mating systems. Indeed, while the *Capsella* genus contains only four species, three are diploid and one is tetraploid. *C. grandiflora* (Klokov) is a diploid and obligate outcrosser (2x=16), with a sporophytic self-incompatibility system, which is geographically restricted to northern Greece and Albania. *C. rubella* Reut. and *C. orientalis* (Fauché & Chaub.) are both selfers (2x=16). *C. rubella* derived from *C grandiflora* ca. 25,000-40,000 years ago (Foxe *et al*., 2009; Guo *et al*., 2009) and its origin seems to be concomitant with the breakdown of the self-incompatibility system (Guo *et al*., 2009). *C. rubella* grows around the Mediterranean and as far north as Belgium in western Europe. The main distribution area of *C. orientalis* extends from central Ukraine to northwestern China and western Mongolia (Hurka *et al*., 2012). *C. bursa-pastoris* (L.) Medik. (Brassicaceae) is an allotetraploid selfing species (4x=32) with disomic inheritance, coming from the hybridization between the ancestors of *C. grandiflora* and *C. orientalis* (ca. 100-300 000 years; Douglas *et al*., 2015). In contrast to its diploid relatives it has an almost worldwide distribution. At least three main genetic clusters can be defined that correspond to: Asia, Europe, and the Middle East. Gene flow among these clusters is low and a strong differentiation both at the nucleotide and gene expression levels was observed (Cornille *et al*., 2016a; Kryvokhyzha *et al*., 2016, 2019b)

Previous studies have investigated differences in competitive abilities in the genus *Capsella* according to ploidy level, mating system and competitors density (Petrone Mendoza *et al*., 2018; Yang *et al*., 2018; Orsucci *et al*., 2020). These studies highlighted clear differences for traits associated to fitness (e.g. number of fruits, germination rate) between *Capsella* species as well as among *C. bursa-pastoris* populations. However, an important limitation of these studies is that they were carried out in growth chambers. Consequently, *C. grandiflora* individuals that are obligate outcrossers did not set seeds because of the absence of pollinators, and flower number instead of fruit or seed sets was used as a fitness proxy. This makes fitness comparison between outcrossing and selfing species difficult and this is why only relative fitness between treatments, instead of absolute fitness, was compared among species. To circumvent this problem, we carried out a common garden experiment within the natural range of *C. grandiflora, C. rubella* and *C. bursa-pastoris*, in northern Greece. We evaluated the sensitivity of the four *Capsella* species to competition under two different environmental, conditions: disturbed (i.e. other plant species are constantly removed) and undisturbed (i.e. other plant species can grow around the focal *Capsella* plant) treatments. We also measured several traits across the plant life cycle. Surprisingly, our experiment showed no significant difference between disturbed and undisturbed treatments. The main differences were actually observed between the three selfers and the outcrosser and unexpectedly ploidy level had limited or no impact on the life history traits measured in this experiment. Overall, *Capsella grandiflora*, the obligate outcrosser, had a higher fitness than all selfing species.

## MATERIAL AND METHODS

### Study material

The experiment comprised 18 accessions of *C. bursa-pastoris* from 18 populations, 10 accessions of *C. rubella* from seven populations, six accessions of *C. orientalis* from four populations and 10 genotypes from 7 different populations of *C. grandiflora*, the outcrossing species (Table 1; details about accessions, see Table S1). The eighteen *C. bursa-pastoris* accessions could further be subdivided into five accessions from the European genetic cluster (EUR), four from the Middle Eastern one (ME), six from the Asian genetic cluster (ASI), and, finally, three from Central Asia (CASI).

**Table 1.**
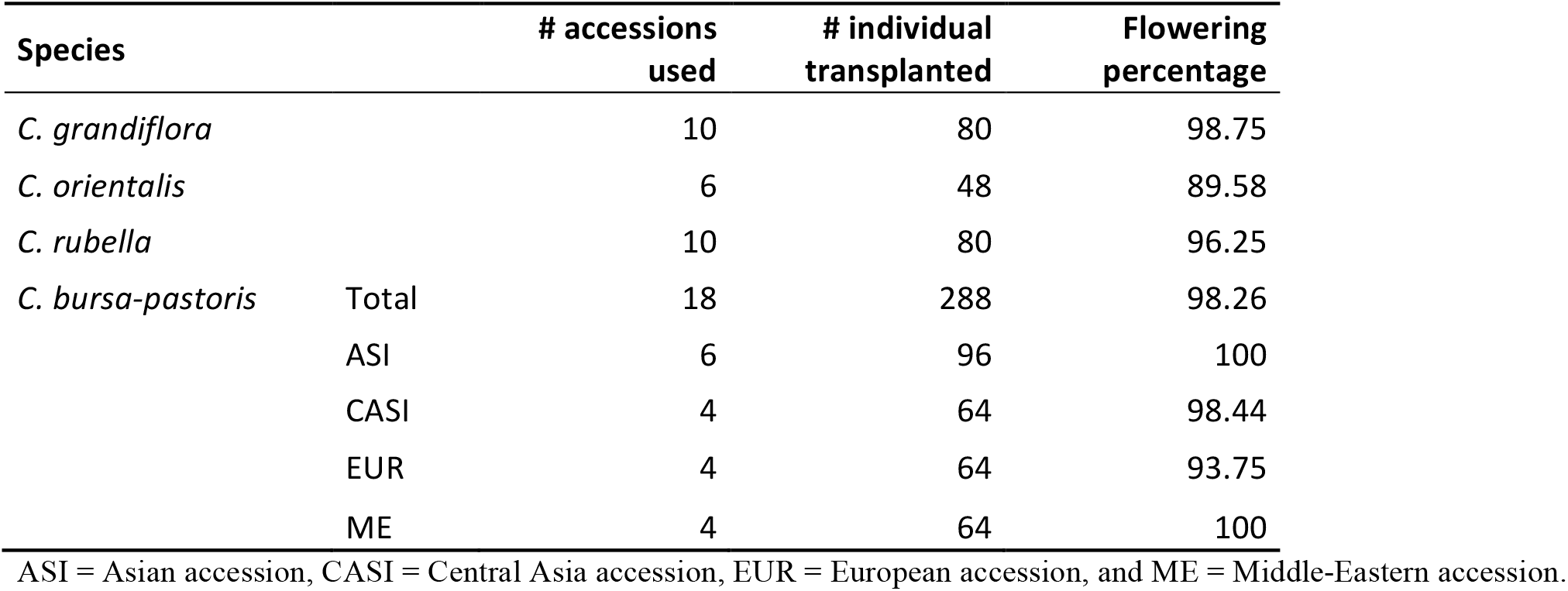
Experimental details and flowering percentage per species and for each geographical origin for *C. bursa-pastoris*

| Species |  | # accessions<br>used | # individual<br>transplanted | Flowering<br>percentage |
| --- | --- | --- | --- | --- |
| *C. grandiflora* |  | 10 | 80 | 98.75 |
| *C. orientalis* |  | 6 | 48 | 89.58 |
| *C. rubella* |  | 10 | 80 | 96.25 |
| *C. bursa-pastoris* | Total | 18 | 288 | 98.26 |
|  | ASI | 6 | 96 | 100 |
|  | CASI | 4 | 64 | 98.44 |
|  | EUR | 4 | 64 | 93.75 |
|  | ME | 4 | 64 | 100 |

ASI = Asian accession, CASI = Central Asia accession, EUR = European accession, and ME = Middle-Eastern accession.

### Experimental design

The experiment was carried out on the campus of the University of Ioannina (Greece) (39°37.090’N and 20°50.806’E), which is located within the natural range of *C. grandiflora*. Differences in competitive abilities between *Capsella* species that differ in ploidy level and mating system were tested under two different treatments: *(i)* disturbed (the vegetation was removed one to two times a week around the *Capsella* focal plant) and *(ii)* undisturbed (without removal of the vegetation).

First, in January 2018 at least 50 seeds per accession were sown into plastic pots (9 x 6.5 cm) containing standard soil (Klasmann Traysubstrat). Pots were first stratified for 7 days by keeping them in the dark at 4°C and then moved into a growth chamber with 16/8h light/darkness cycles at 22°C to allow germination. Germination of the earliest accessions started approximately one week later (Table S1). Seedlings that presented two pairs of well-developed first true leaves were transplanted in single square pots (8 x 8 cm). In each pot two seedlings were planted on opposite corners. At the beginning of March 2018, all pots were moved to an experimental garden for a phase of acclimation to external conditions.

Two weeks before transplantation in the experimental plots, vegetation and larger stones were removed from the experimental field, thereby homogenizing the topsoil. The experimental field was then covered with anti-weed tissue, in which 10 x 10 cm holes separated by 20 cm were done to create experimental plots. To facilitate transplantation, vegetation was removed in all plots just before performing the transplants. Then, vegetation was removed once to twice a week in plots corresponding to the disturbed treatment, while plots corresponding to the undisturbed treatment were left untouched.

From the 7^th^ to the 14^th^ of April, a total of 496 plants were distributed randomly (*sample* function, R version 3.6.3; R Development Core Team, 2020) over the two different treatments which were organized in four experimental groups. The 496 plants corresponded to eight replicates of eighteen accessions of *C. bursa-pastoris*, four replicates of ten accessions of *C. grandiflora*, four replicates of ten accessions of *C. rubella* and four replicates of six accessions of *C. orientalis*. In addition, in each treatment, 20 plots (also distributed randomly) were left empty in order to monitor spontaneous vegetation recolonization.

The performance of the four *Capsella* species under the two different treatments (disturbed vs. undisturbed) was monitored by recording for each plant: *(i)* vegetative traits such as rosette’s diameter (measured on transplantation’s day) and number of floral stems by plants (measured on dead plants), *(ii)* phenological traits such as flowering start (*i*.*e*. number of days between plant transplantation and the first flower) and lifespan (*i*.*e*. number of days between plant transplantation and death) and *(iii)* main fitness components, i.e. fertility measured as the number of fruits, which is a proxy of number of offspring and viability quantified by seeds germination rate. In addition, we computed a relative fitness index (*W*_*index*_), which is an integrative measure of overall performance and is defined as the product of number of fruits and germination rate. The combination of these two components provides a better estimate of the overall fitness of each species (and genetic cluster within *Cbp* species). This index varies between 0 and +∞; *W*_*index*_ = 0 means that the fitness of the individual is low (due to a low performance of at least one of its components) and a large value of *W*_*index*_ indicates that the individual performed well for both fitness components.

In addition to these traits, damages (cut stems, unhealthy plant, etc …) on the plants, which could be caused by phytophagous insects (ants, aphids) or water stress, were recorded and synthetized in a unique variable called *damages* (see below).

For all four species, seeds were collected on each flowering plant to quantify the germination rate. One year after the common garden experiment was over, 50 randomly sampled seeds per plant were sown directly in soil of square pots (4 x 4 cm) that were randomized. After ten days of stratification (24h dark, 4°C), the pots were moved into growth chambers (12:12h light:dark, 22°C). The number of seedlings after 21 days was recorded and the germination rate was calculated as the number of seedlings over the number of seeds sown.

### Statistical analysis

All statistical analyses were performed using the R software (R version 3.6.3; R Development Core Team, 2020). First, in order to evaluate the contributions of each trait to the variation among species, principal component analyses were performed on all traits with the function *dudi*.*pca* (ade4 package; (Dray & Dufour, 2007). The resulting PCs were then used to limit redundancy and allow a reduction of the number of variables investigated in further analyses; the traits *Ants, Aphids, Stress* and *Inflorescence damage* (Fig 1A) were highly correlated and were merged into a unique variable called *damages*. Because *number of fruits* and *number of stems* were also highly correlated (Pearson’s product moment correlation: t = 16.015, df = 494, r =0 .6, *p* < 0.001, Fig. 1A) we only used *number of fruits* in all analyses.

**Figure 1.**
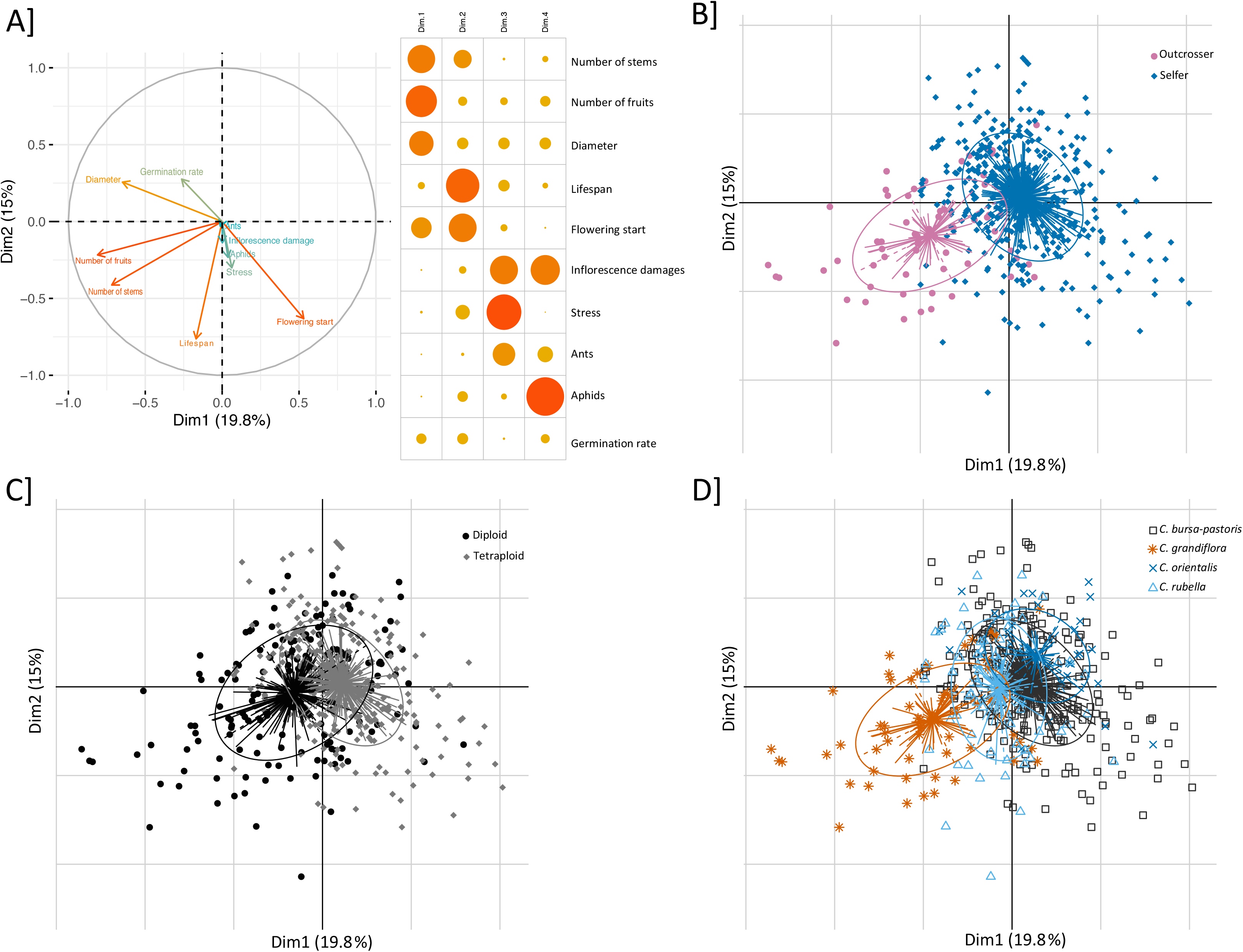
Principal component analysis (PCA) based on life history traits measured in *Capsella* spp. A]. Correlation circle (left panel) showing the relative contribution of each variable to the two first principal components explaining 35% of the total observed variance; the right panel shows the relative contribution of each variable to the four first principal components (the darker and the larger the discs the higher the contribution). **B-C-D]** PCA with all the individuals using different grouping to highlight differences relative to mating system (**B**), ploidy level (**C**) and species (**D**).

Then, for all traits (*diameter* of the rosette, *lifespan, flowering start, number of fruits, germination rate* and *fitness index*), linear models were used to test the effect of four main biological characteristics: (i) the reproductive system (outcrossing *vs* selfing), (ii) the ploidy level when focusing on selfing species (diploid vs tetraploid), (iii) the species or (iv) the genetic cluster, when focusing on *C. bursa-pastoris*.

For a given trait, the fixed predictors were (i) the treatment (qualitative variable, two levels: disturbed or undisturbed), (ii) the biological characteristics (qualitative variables, two levels for reproductive system and ploidy and four levels for species and genetical clusters of *Cbp*, see above for more details) and (iii) the interaction between the two predictors. We added one co-variable, which measures damages (qualitative variable, two levels: damaged or not). In addition, a row effect (which is due to experimental constraints) and an accession effect were included as random effects in each model.

We analysed the rosette size (quantitative variable) and the flowering start (LOG10 normalization) with a linear model, assuming a Gaussian error distribution (*lmer* function, stats package, R version 3.6.3; R Development Core Team, 2020). Lifespan, Number of fruits, fitness index and germination rate were analyzed using a generalized linear mixed model, assuming a negative binomial distribution, except for the last one where we assumed a binomial distribution (*glmer*.*nb* and *glmer* functions, respectively used, lme4 package, Bates *et al*., 2015).

The significance of the fixed-effects was tested with deviance analyses using the Anova function of the car package in R (Fox & Weisberg, 2019). Interactions were tested first (type III ANOVA), then, if the interaction was not significant, a model without interaction term was run to test only the main fixed effects (type II ANOVA). Significance of difference between factor-levels was tested by a post-hoc procedure allowing multiple comparisons (*glht* function, multcomp package, Hothorn *et al*., 2008).

## RESULTS

The two first principal components of the PCA including all traits explained 35% of the total observed variance (Fig. 1A) and the variables with the highest contributions were the number of fruits and stems, the rosette diameter and the phenological traits. More specifically, the first principal component discriminated the two reproductive systems (Fig. 1B): PC1 separated *C. grandiflora*, which is the only outcrosser, from the three selfing species (Fig 1.D) (selfer *vs* outcrosser; Fig. 1B) and, to a lesser extent, the two ploidy levels (diploid *vs* tetraploid, Fig 1.C).

As the damage-related variables (ants, aphids, short water stress or cut inflorescence) had a very small contribution to the proportion of variance explained and since their contributions were of similar magnitude (Fig. 1A), they were grouped into a single variable: *damages* (see Materials and Methods section “Statistical analyses”).

### Absence of treatment effect

Unexpectedly, no differences between disturbed and undisturbed treatments were observed for most of the response variables (Fig. S1, Table 2). Thus, the presence of other species around the focal *Capsella* individuals did not affect their development (e.g. rosette size), life cycle (phenology) and fertility. Surprisingly, seeds from the plants that grew up in disturbed treatments, i.e. in plots in which the vegetation was removed, tended to have a higher germination rate than those from the plants that grew up in a treatment with competitors (Fig 2B). It is also noteworthy that germination rate is the only trait for which the interaction between individuals and their environment was significant (Table 2). Finally, while no direct effect of the environment on the plant was observed, the environment could still impact the next generation by decreasing the germination success of the offspring of plants exposed to a stronger competition (Fig 2B, Table 2)

**Table 2.**
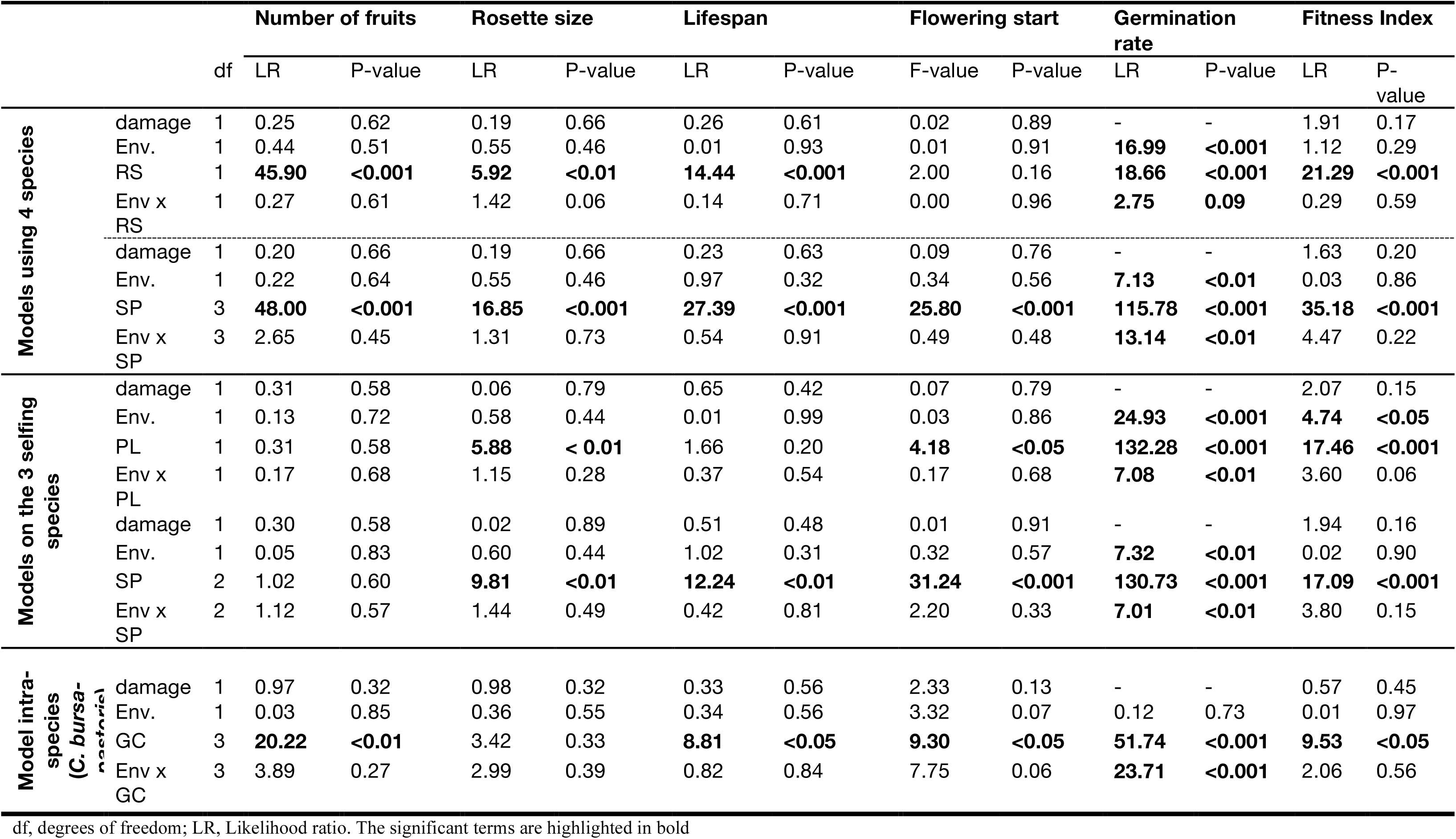
Analysis of deviance for all the measured traits and the Fitness Index

**Figure 2.**
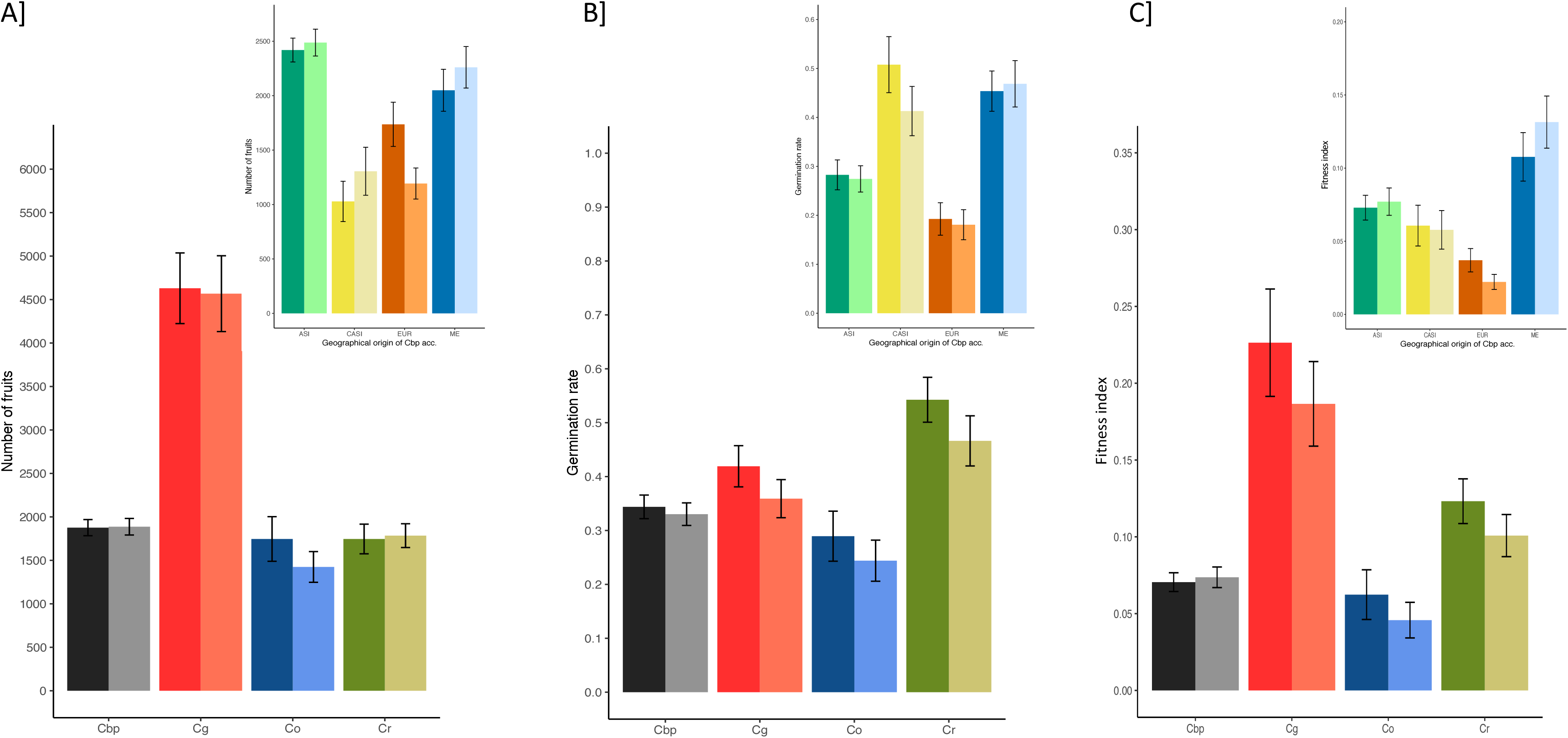
Life history traits and fitness index for *Capsella* species (main figure) and between genetical clusters of *C. bursa-pastoris* (top right panels). The mean number of fruits (**A**), the mean germination rate (**B**) and the mean fitness index (**C**) which is scaled to vary between 0 and 1 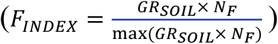 meaning that the higher the index, the higher the relative fitness is. The three traits are represented for *C. bursa-pastoris* (Cbp, in gray), *C. grandiflora* (Cg, in red), *C. orientalis* (Co, in blue) and *C. rubella* (Cr, in green) or genetical clusters of *C. bursa pastoris* (ASI in green, CASI in yellow, EUR in orange and ME in blue) in disturbed (dark color) and undisturbed (light color) treatments. The corresponding standard errors are indicated by the bars.

### Traits were primarily influenced by reproductive system

Except for *flowering start*, the outcrosser species, *C. grandiflora*, differed from selfing species for all other traits. First, the rosette diameter was significantly larger in outcrossing individuals (c.a. 12.3 ± *2*.*6* cm) than in selfing ones (10.6 ± 3.2 cm; Tables 2 and 3). Second, the outcrossers produced about 2.5 times more fruits than selfers (out. = 4668, self. = 1884; Table 2 and Fig. 1A). This increase in term of fruits production could be assigned to the extended lifespan of the outcrossers (out. = 130, self. = 118 days; df = 474, z = −6.92, *p* < 0.001; Tables 2 and 3). More precisely, since flowering start did not differ significantly between the two groups (out. = 16, self. = 17 days; df = 474, t = 0.219, *p* = 0.83; Table 3), the increase in fruits production is due to an extended reproductive period in outcrossers (Fig. S2). Finally, germination rate was also on average higher among outcrossing accessions than among self-fertilizing ones (out. = 0.39, self. = 0.35; Table 2 and Fig. 1B).

**Table 3.**
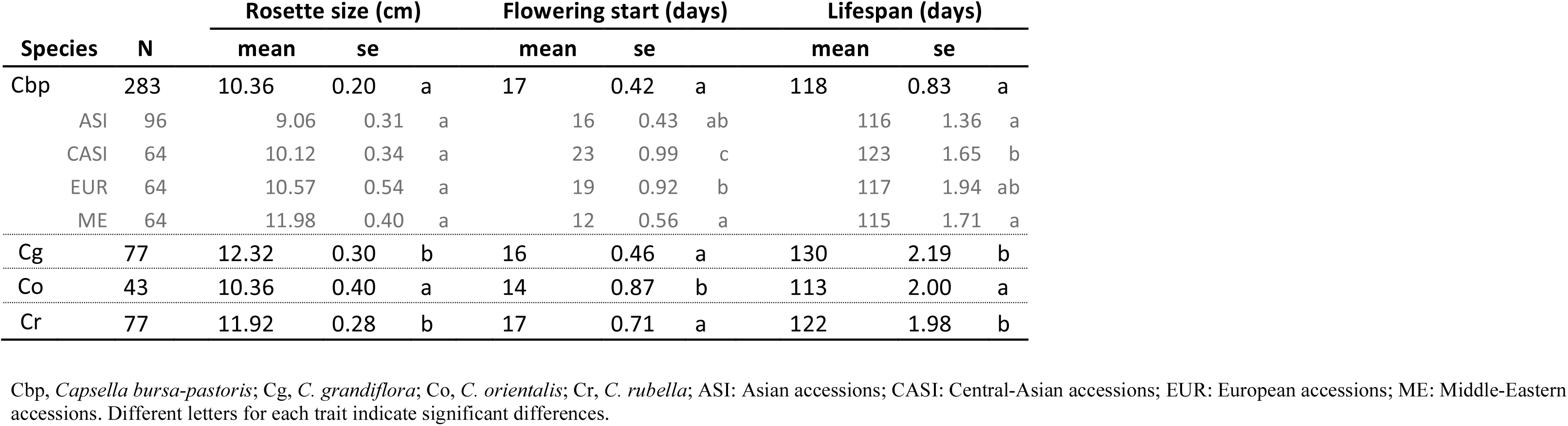
Raw data for traits indirectly linked to the fitness for the four species (in black) and within Cbp according the geographical origin (in gray)

### Weak effect of polyploidization

Because of the strong effect of the mating system, we limited the analysis of the effect of ploidy to the three selfing species. Surprisingly, ploidy had a weak, and generally non-significant, effect on most traits (Table 2). However, its effect on both rosette size and germination rate, and consequently also on *W*_*index*_, was significant: (i) rosette size was slightly smaller in *C. bursa-pastoris*, (diameter = 10.2 ± *3*.*4* cm) than in *C. rubella and C. orientalis*, (diameter = 11.1 ± *3*.*1* cm; Tables 2 and 3), and (ii) germination rate and fitness index were lower for the tetraploid species than for the diploid ones. On average flowering also started later in the tetraploid species than in the two diploid ones. However, these results are largely due to extreme values occurring for one or the other of the diploid species and therefore are not really associated to ploidy level *per se* but instead to a species effect (Table 2 and Fig. 2 B and C). This is well illustrated by germination rate that is higher in *C. rubella* than in both *C. bursa-pastoris* and *C. orientalis*, or flowering time which occurs earlier in *C. orientalis* than in *C. bursa-pastoris* and *C. rubella*.

### Foreign individuals differ phenotypically from local ones

Phenotypic traits were strongly affected by the geographical origin of the plants. Indeed, the second principal component, which explained 16% of the total variance (Fig. 1A), mainly discriminate the samples according to their geographical origin (Fig S3). Within *C. bursa-pastoris*, there was also clustering according to local geographical origin (genetic clusters, Fig S3).

*C. orientalis* was as an outlier which is characterized by *(i)* a smaller rosette size, *(ii)* a shorter lifespan and *(iii)* an earlier flowering start (Table 3). *C. orientalis* is the only species that does not occur naturally in Greece. This pattern could be due to local adaptation to environmental conditions since individuals of *C. bursa-pastoris* from Asia and the Middle-East also lived shorter and flowered earlier (Table 3) than European and Central Asian accessions. They bore more fruits (Fig 2A) too. However, the grouping was different for germination as Asian and European accessions germinated significantly less than accessions from the Middle-East and Central-Asia (Fig 2B).

### Lifetime reproductive success: towards an integrative measure incorporating major fitness components

The fitness index magnified the differences between the four species as well as between the four genetic clusters within *C. bursa-pastoris* (Fig. 2C). As expected, when number of fruits and germination rate are jointly considered, selfing species have a lower fitness than *C. grandiflora* (Table 2; Fig. 2C). Among the selfing species, the fitness index also showed that *C. rubella* globally performed better than *C. bursa-pastoris* (df = 444, z = 3.364, *p* < 0.01; Fig. 2C) and *C. orientalis* (df = 444, z = 3. 138, *p* < 0.01; Fig. 2C) the last two species being similar (df = 444, z = −1.322, *p* = 0.53; Fig. 2C). Such a result was not visible when the two traits were considered separately (Fig 2. A and B). Considering the fitness index also changed the ranking among the four genetic clusters in *C. bursa-pastoris:* while Asian accessions produced a significantly higher number of fruits than plants from the three other genetic clusters they had a lower overall performances than Middle East accessions when germination rate was also considered (df = 265, z = 3.408, *p* < 0.01; Fig. 2).

## DISCUSSION

Plants of the four extent *Capsella* species were grown in disturbed and undisturbed environments and their phenotypes and lifetime reproductive success evaluated. To the best of our knowledge, this is the first time *C. grandiflora* is compared to its self-fertilizing relatives under conditions where it can be pollinated and therefore the comparison is based on direct estimates of fitness components. Surprisingly, the environmental disturbance used here did not lead to fitness differences among species. Such a result was unexpected because the effect of competition had been demonstrated on the same species in several experiments conducted under controlled environments (Petrone Mendoza *et al*., 2018; Yang *et al*., 2018; Orsucci *et al*., 2020). In previous greenhouse experiments, the density of competitors was adjusted to ensure detecting significant competition effects. Here, the natural environment appeared less competitive that we thought. Ploidy had a weak positive effect on rosette size, flowering start and germination rate but this was principally due to the low performance of *C. orientalis*, the only species not naturally found in sympatry with *C. grandiflora*. However, both mating system and geographical origin had a strong effect on fitness components. The lack of effect of competition contrasts with previous studies carried out in growth chambers (Petrone Mendoza *et al*., 2018; Yang *et al*., 2018; Orsucci *et al*., 2020). A strong difference in flower (not fruit) number was observed between selfers and outcrossers in previous experiments in growth chambers but was not interpreted because it could have been driven by the absence of pollinator inducing on-going flower production in the outcrossing species (Petrone Mendoza *et al*., 2018; Yang *et al*., 2018). Similarly, absolute fitness proxies were not compared between diploid and tetraploid selfers but the difference in number of flowers was small, the two diploids producing only slightly more flowers than the tetraploid. The current results suggest that, despite artificial conditions, the fitness measures in greenhouse were rather good proxies of difference among species in natural conditions. Below we discuss some of the caveats of the experiment and the practical implications and evolutionary consequences of the results.

### Conducting experiment under semi-natural conditions leads to different outcomes than under controlled conditions

Biological systems are complex and dynamic systems and they depend on multiple biotic and abiotic factors and their interactions. Consequently, to understand the effect of specific factors on a response variable, it is often necessary to resort to experiments under controlled conditions. A first experiment under controlled conditions by Petrone *et al*. (2018) aimed to understand the effects of (i) mating system, (ii) ploidy level and (iii) their interaction on direct and indirect components of fitness. In Petrone *et al*. (2018) both ploidy and mating system influenced the response to competition: diploid selfers (*C. rubella*) were more sensitive to competition than diploid outcrossers (*C. grandiflora*), and tetraploid selfers (*C. bursa pastoris)*, which did not differ significantly. However, the outcrosser, alone or in presence of competitors, produced more flowers than either selfing species, showing that the outcrosser had a higher absolute fitness. However, because of the absence of pollinators the results were probably biased in favour of outcrossers: since outcrossing individuals did not need to invest energy and resources in seed development, they could produce more flowers and live longer. A main goal of the present study was to confirm the results obtained by Petrone *et al*. (2018) under semi-natural conditions, in the native geographical range of *C. grandiflora*, where pollinators are present and where plants will face natural competitors/enemies and climatic conditions. We confirmed that the fertility of the outcrossing species was higher than that of the three self-fertilizing species. While we obtained a similar ranking between species (Table S2), we showed that conducting the experiment under semi-natural condition increased significantly the number of fruit produced (*C. bursa pastoris*: t = 19.568, df = 369; *C. grandiflora*: t = 11.789, df = 83.628; *C. rubella*: t = 10.361, df = 88.162; all *p-value* < 0.001). However, it should be noted that the number of fertile fruits may have been overestimated due to our counting method. We estimated the number of fruits by counting the number of pedicels on each individual at the end of flowering, but we could not differentiate between fully developed fruits and non-fertilized fruits because a large number of fruits were already opened and had shed seeds. Although the absolute value of the number of fruits may have been overestimated, the comparison between species is still valid since the counting method is the same for all species. Regarding polyploidy, the hypothesis that doubling the genome could help alleviate problems associated to self-fertilization has not been verified in this study. In some analyses, the tetraploid selfer performance was even worse than the performance of diploid selfers, but this result was mainly due to the higher performances of *C. rubella* in comparison with *C. bursa-pastoris* and *C. orientalis*, the latter two having the same pattern for most of the traits measured. In order to have a real idea of the effect of polyploidization, studies based on a larger number of diploid and polyploid species, for instance along the Brassicaceae lineage, would be needed.

Unlike previous studies conducted under controlled conditions on these species (Petrone Mendoza *et al*., 2018; Yang *et al*., 2018; Orsucci *et al*., 2020) response to competition could not be assessed in the present study simply because there were no differences in response to the two treatments to which the plants were exposed (disturbed and undisturbed). Several, non-mutually exclusive, factors could explain why the fitness trait (*i*.*e*. the number of fruits) was similar between disturbed and undisturbed plots. First, the two types of plot were actually disturbed to permit the transplantation of *Capsella* individuals and as the four species used in this experiment are ruderal species (*i*.*e*. plant species that are first to colonize disturbed environments), their establishment was facilitated compared to potential local competitors. Second, the plots were too close to each other and all plants were affected by competition to the same extent at the root level. Third, the effect of the competitors was negligible because the resources in the field were not very limiting compared to the resources available in the laboratory (ground vs. soil in a pot).

### Possible confounding effect of local maladaptation

In our study, we found a strong effect of geographic origin: individuals coming from plants that have been sampled in Greece or around the Mediterranean area (*i*.*e*. all accessions of *C. rubella* and *C. grandiflora* and the accessions corresponding to ME genetic cluster of *C. bursa-pastoris*) had a higher fitness index than foreign genotypes (*i*.*e*. all accessions of *C. orientalis*, and the accessions corresponding to ASI, CASI and EUR genetic cluster of *C. bursa-pastoris*). These genotypes could be maladapted to the local conditions of the experiment, which could indirectly contribute to differentiation among selfing species and populations. Indeed, plant populations are often found to be locally adapted (Savolainen *et al*., 2007; Hereford, 2009). However, little evidence of local adaptation was detected in *C. bursa-pastoris*, at least at large geographical scale (Cornille et al., unpublished): ASI populations have been shown to perform worse than other populations in several environments, including in their native one in China (Cornille *et al*., 2016b), in agreement with their higher genetic load (Kryvokhyzha *et al*., 2019b).

### Phenological, vegetative and fitness-linked traits are strongly differentiated in the outcrossing species

The strongest differences for both life-cycle traits and fitness components were between selfing and outcrossing species. In particular, *C. grandiflora* produced 2.5 times more flowers, had a reproductive period on average 14 days longer and a rosette size about 2 cm larger than the selfing species. These results do not match the phylogenetic relationships among the four species. Instead, they support strong trait convergence between selfing diploid species that diverged at very different time from *C. grandiflora*: *C. orientalis* diverged around one million years ago while *C. rubella* diverged only 20,000 to 50,000 years ago (Wozniak et al. 2020). Similarly, in the tetraploid, while parental legacy is relatively well balanced at the expression level (Kryvokhyzha *et al*., 2019a), morphological traits are clearly biased towards the selfing parental species. This suggests that evolution of selfing not only affects floral morphology but also many other traits, pointing to a more global and integrated selfing syndrome including both vegetative and ecological traits (see discussion in Petrone Mendoza *et al*., 2018). This is in line with the time-limitation hypothesis that posits that selfing annuals plants invest less in developmental traits (decreasing flower and plant sizes) and have shorter bud development time and flower longevity than outcrossing relatives (Snell & Aarssen, 2005). This hypothesis, which predicts that selfing annuals plants are under strong *r*-selection (i.e. the species bet on reproduction with a high rate of growth to compensate for low chance of survival) and, by consequence, that the period to complete their life cycles is severely limited, could be involved in these differences between selfers and outcrossers, especially for species found in disturbed habitats, like roadsides or cultivated fields (Aarssen, 2000; Snell & Aarssen, 2005). In addition, selfing species often exhibit a higher genetic load than their outcrossing relatives (Glémin & Galtier, 2012; Arunkumar *et al*., 2015). This is the case in the *Capsella* genus (Kryvokhyzha *et al*., 2019b) and it could also contribute to lower their fitness.

Selfing confers a reproductive assurance advantage over outcrossing and thereby facilitates colonization of new territories. In the *Capsella* genus, all selfing species had a larger geographical range than their outcrossing relative, *C. grandiflora*, which has a very limited distribution range, being confined to a small geographical region in northern Greece and Albania. This limited geographical range of *C. grandiflora* is, at first glance, surprising considering that it seems to perform better than all its selfing relatives, has a higher genetic diversity (Kryvokhyzha *et al*., 2019b) and a better competitive ability than diploid selfers (Petrone Mendoza *et al*., 2018; Yang *et al*., 2018). Actually, the factors limiting its spread are still unclear but it does not seem to be due to a dependence on specific pollinators (pers. observations and communications). A potential explanation of its limited range could be a low diversity at self-incompatibility alleles driving to few fertile and viable crosses in marginal populations, hitherto undiscovered ecological factors or human activities altering its natural habitat, a disruption between plant and pollinators limiting the gene flow and species expansion (*i*.*e*. fragmented environment could be responsible, in part, of the success of selfing reproductive systems; Eckert *et al*., 2009). Finally, the above line of arguments assumes that all *Capsella* species are strict annuals, such that fitness can be compared among species and population through a single round of reproduction. However, shorter life cycle and quicker development may allow a second reproductive episode in selfing species. If so, differences in cumulative fitness over one year between selfers and outcrossers should differ less than anticipated. In particular, flowering individuals of *C. bursa-pastoris* can be observed almost all over the year (pers. obs.), and we still do not know whether it corresponds to several generations or to sparse germination from a seed bank across a given year. In any case, further studies will be needed to find the causes to the limited geographical range of *C. grandiflora*.

### Importance to quantify different life history traits throughout lifespan to get a more accurate picture of long-term reproductive success

Fitness can be estimated in many different ways; however, it has often been estimated by only one of its components, for example the number of seeds produced in plants (Primack and Kang 1989). We note that this is problematic since fitness depends on multiple components and is tightly associated to the environment under which it is measured. Often, fitness components are analysed separately because each component is more conform to a simple parametric distribution than fitness measured over the lifespan (Shaw *et al*., 2008). However, the fitness of an individual depends on its ability to survive, to find a mate, to produce viable and fertile offspring, and a minimal definition of fitness would then be Fitness = Survival × Fecundity. Short of that, one is explicitly or implicitly making some crucial assumptions. For example, when one assumes that plants that produce the highest number of seeds have the highest fitness, one implicitly also assumes that there is no trade-off between seed number and seed quality. However, a seed number/seed size trade-off is actually expected if the resources to be invested in reproduction are limited. Under a simple model of resource allocation, the distribution of limited resources among several seeds involves a reduction of the amount of resources invested in each seed with an increase of seed number (Smith & Fretwell 1974, Shipley & Dion 1992). A lower investment of resources in seeds could, in turn, affects seeds germination, growth, survival but also fecundity of adult plants of the next generation (Eriksson 1999, Tremayne et al. 2000, Lehtilä & Ehrlén 2005). Hence, the presence of trade-off between traits shows that equating fitness to a single of its component at a single time point of the life cycle can severely alters the biological interpretation of the results.

In the present study, the global trends shown by the two components of fitness *number of fruits* and *germination rate*,with respect to the impact of mating system, were similar to those observed for the fitness index which integrates both fertility and viability: namely, the overall performance of the outcrossing species was higher to those of the selfing species. However, the relative performances among selfing species changed, with *C. rubella*, a Mediterranean species, globally having a higher fitness index than the cosmopolitan *C. bursa-pastoris* and the Asian *C. orientalis*. Hence, taking into account at least two direct components of fitness allowed us to reach more robust and meaningful conclusions than the analysis of a single trait or each trait independently would have done. However, it should not be forgotten that other components of fitness may also matter and that they are generally highly variables, such as length and timing of the flowering season, resistance to pests and diseases or plant size (Brown 1975 cited by Primack and Kang 1989).

## Conclusion

Although we did not detect any differences between disturbed and undisturbed treatments, the present study highlighted how profound the impact of the shift in mating system is. Indeed, self-fertilizing individuals had lower performances than cross-fertilizing ones. We also showed that a possible local adaptation pattern also amplified differences among plants within and between species. Finally, the present study further demonstrated the importance of taking into account several life history traits at different times across the life cycle to reach a better estimate of fitness.

## DATA ACCESSIBILITY

The datasets supporting this article will be uploaded in Dryad repository.

## AUTHOR’S CONTRIBUTIONS

MO, MG, ML and SG conceived the study. TV, MG and TD prepared the experimental field; TV managed the plants growth, the monitoring of the experiment and the data collection; MO performed the germination test in growth chamber; MO performed the statistical analyses. MO and ML wrote the manuscript with input from other authors.

## ACKNOWLEDGEMENTS

We thank P. Milesi for constructive discussions and comments on the manuscript, George Chioureas for his help with fieldwork and Hera Karayanni for hosting us in her laboratory at the University of Ioannina.

---

**Figure S1.**
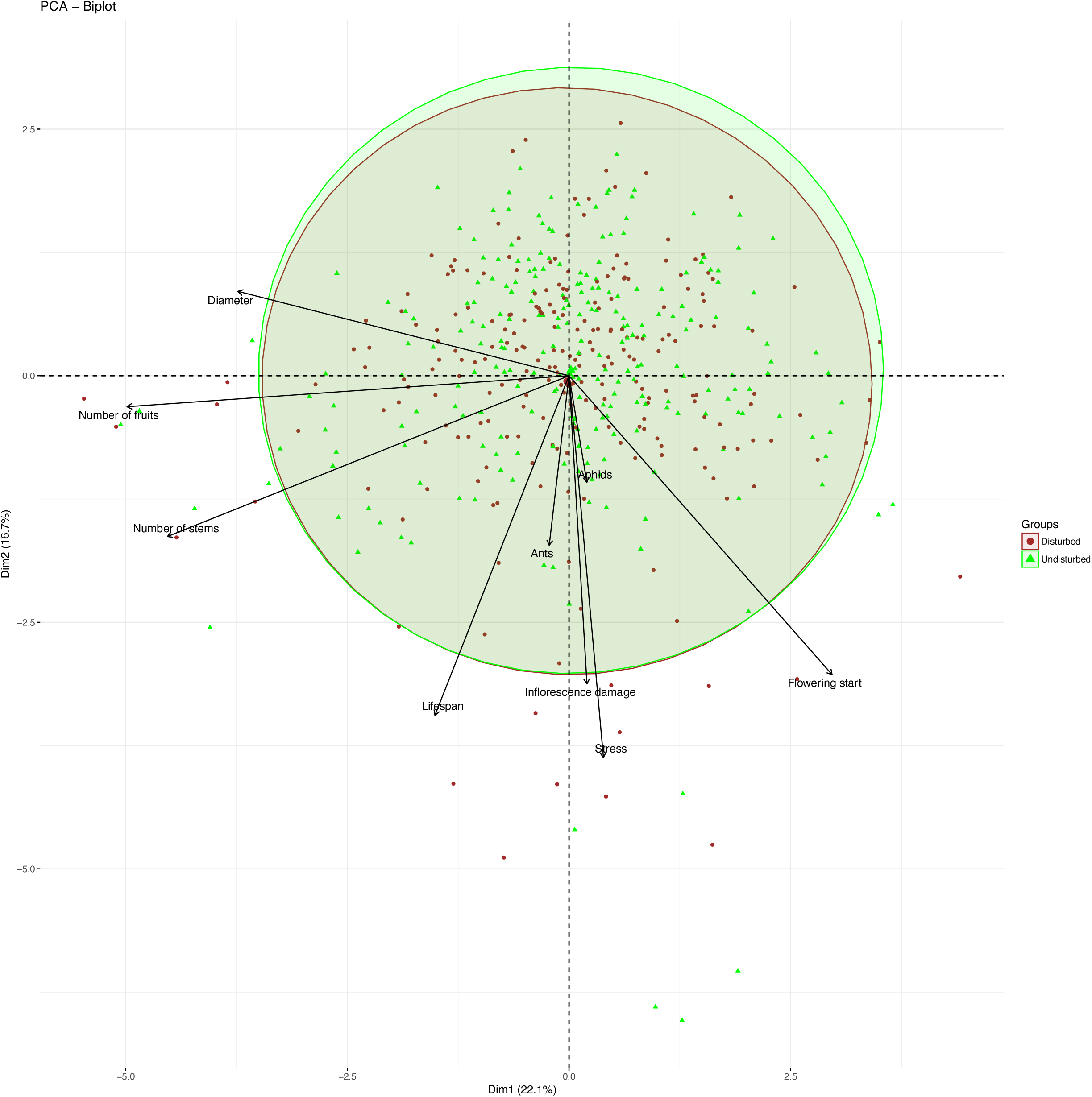
Principal component analysis (PCA) based on life history traits measured to highlight differences relative to the treatment. Individuals from disturbed (red dots) and undisturbed (green triangle) treatments are represented whatever the species. The relative contribution of each variable to the two first principal components is proportional to the length of the arrow (the longer the arrow, the more the variable contributed to the explained variance).

**Figure S2.**
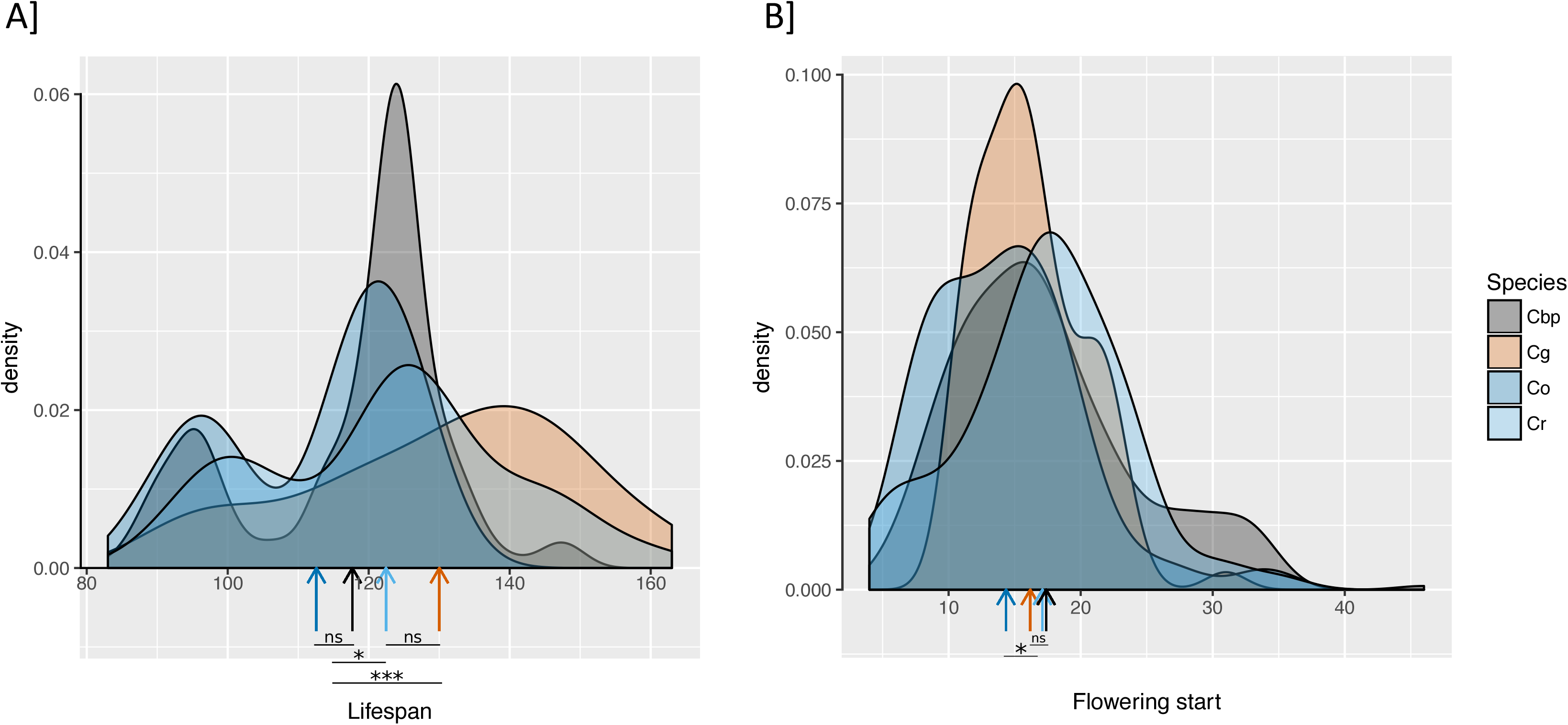
Phenological differences between *Capsella* species. Lifespan (A) and flowering start (B) measured as the number of days after transplant, are represented for *C. bursa-pastoris* (Cbp, in gray), *C. grandiflora* (Cg, in red), *C. orientalis* (Co, in dark blue) and *C. rubella* (Cr, in light blue). The vertical arrows on the x-axis indicate the mean lifespan (A) and the mean flowering start for each species. The stars indicate significant differences between the four species (*p* < 0.05 *, *p* < 0.01 **, *p* < 0.001 ***).

**Figure S3.**
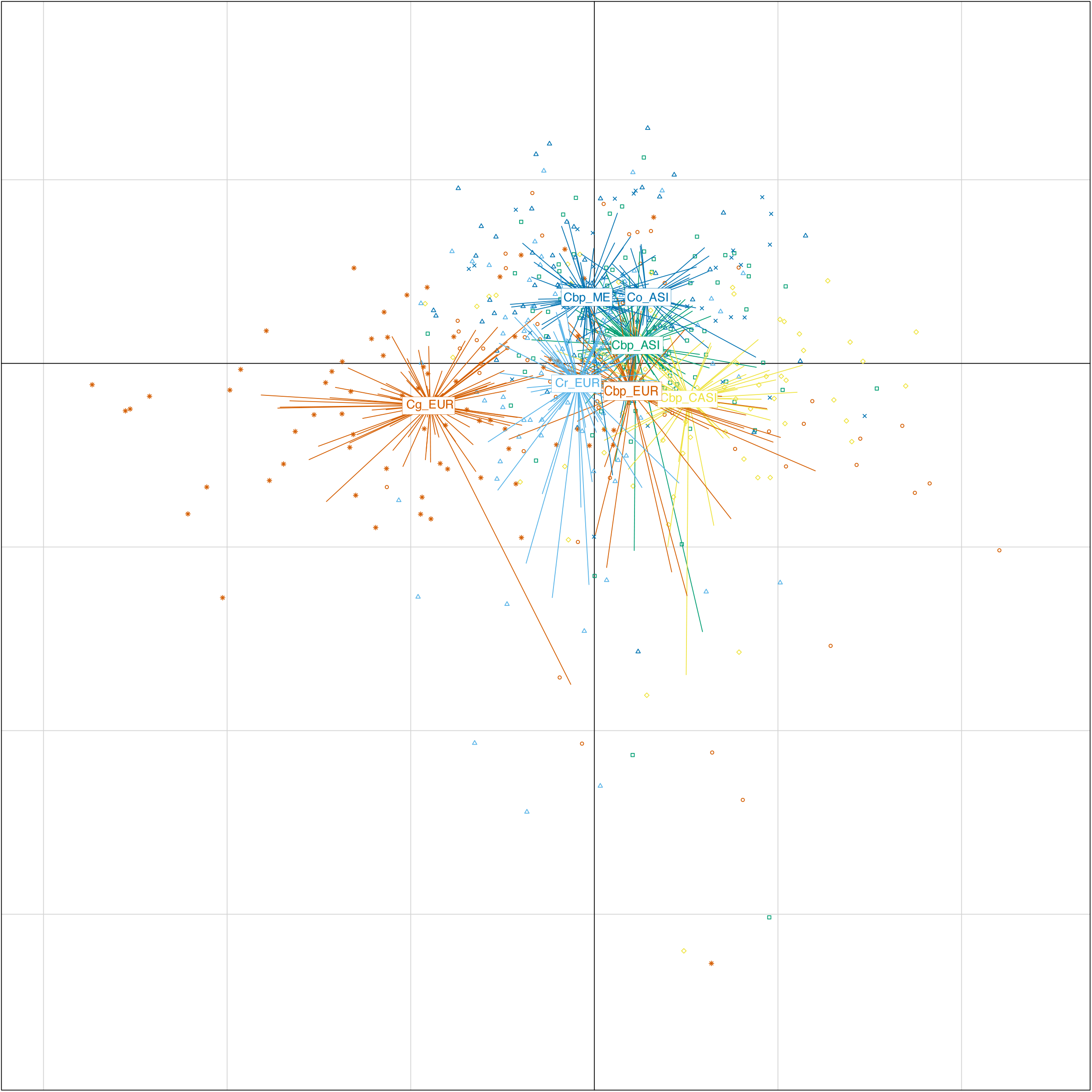
Principal component analysis (PCA) based on life history traits measured to highlight the different genetical clusters within *Capsella bursa pastoris* : European cluster (Cbp_EUR, orange dot), Asian cluster (Cbp_ASI, green square), Middle-Eastern cluster (Cbp_ME, dark blue triangle) and Central Asian cluster (Cbp_CASI, yellow diamond) and showing their relative position to the three other *Capsella* species: *C. grandiflora* (Cg_EUR, orange star), *C. orientalis* (Co, dark blue crosses) and *C. rubella* (Cr, light blue triangle).

**Table S1.** Sample information

| Species | Accession ID | Accession name | Origin | Latitude | Longitude | Germination date (> 10 seeds) |
| --- | --- | --- | --- | --- | --- | --- |
| *C. bursa pastoris* | Cbp2 | SE33 | EUR | 56.15 | 13.77 | 25/02/2018 |
|  | Cbp8 | VLA3 | EUR | 43.13 | 131.92 | 24/01/2018 |
|  | Cbp9 | IRRU2 | EUR | 52.24 | 104.27 | 23/01/2018 |
|  | Cbp10 | HRB132 | EUR | 45.45 | 126.37 | 23/01/2018 |
|  | Cbp11 | TR73 | ME | 41.02 | 28.97 | 22/01/2018 |
|  | Cbp12 | JO56 | ME | 31.97 | 35.98 | 22/01/2018 |
|  | Cbp13 | AL87 | ME | 35.45 | 7.96 | 22/01/2018 |
|  | Cbp14 | WAC5 | ME | 31.55 | -97.15 | 22/01/2018 |
|  | Cbp15 | PL | ASI | 23.99 | 120.96 | 20/02/2018 |
|  | Cbp18 | HJC421 | ASI | 30.16 | 120.13 | 20/02/2018 |
|  | Cbp20 | JZH152 | ASI | 30.2 | 112.06 | 24/01/2018 |
|  | Cbp21 | HY86 | ASI | 31.76 | 35.17 | 26/02/2018 |
|  | Cbp22 | XN444 | ASI | 36.37 | 101.46 | 20/02/2018 |
|  | Cbp23 | TBS195 | ASI | 33.57 | 107.45 | 15/02/2018 |
|  | Cbp27 | KIRG-5-12 | CASI | 39.79 | 72.18 | 24/01/2018 |
|  | Cbp29 | PETER-RUS-10 | EUR | 59.93 | 30.33 | 20/02/2018 |
|  | Cbp30 | TACH-CHIN-6 | CASI | 47.07 | 83.01 | 23/01/2018 |
|  | Cbp31 | BERG-CHIN-3 | CASI | 48.33 | 87.12 | 23/01/2018 |
| *C. orientalis* | Co1 | QH-CHIN-1 | ASI | 46.70 | 90.83 | 23/01/2018 |
|  | Co5 | URAL-RUS-6 | CASI | 55.11 | 61.39 | 23/01/2018 |
|  | Co6 | URAL-RUS-18 | CASI | 55.11 | 61.39 | 23/01/2018 |
|  | Co7 | GUB-RUS-2 | CASI | 51.29 | 58.18 | 23/01/2018 |
|  | Co9 | OKT-RUS-2 | CASI | 54.32 | 62.68 | 23/01/2018 |
|  | Co10 | OKT-RUS-8 | CASI | 54.32 | 62.68 | 23/01/2018 |
| *C. grandiflora* | Cg1 | Cg1.1 | EUR | 39.66 | 20.85 | 28/01/2018 |
|  | Cg14 | Cg1.37 | EUR | 39.66 | 20.85 | 28/01/2018 |
|  | Cg16 | Cg2.19 | EUR | 39.88 | 20.71 | 28/01/2018 |
|  | Cg17 | Cg2.2 | EUR | 39.88 | 20.71 | 24/01/2018 |
|  | Cg27 | Cg5.27 | EUR | 39.90 | 20.71 | 25/01/2018 |
|  | Cg33 | Cg57 | EUR | 39.96 | 20.71 | 20/02/2018 |
|  | Cg35 | Cg26 | EUR | 39.30 | 20.98 | 25/01/2018 |
|  | Cg36 | Cg3 | EUR | 39.41 | 20.83 | 30/01/2018 |
|  | Cg37 | Cg9.1 | EUR | 39.41 | 20.83 | 23/01/2018 |
|  | Cg40 | Cg9.2 | EUR | 39.96 | 20.71 | 25/01/2018 |
| *C. rubella* | Cr1 | Cr1.1 | EUR | 41.69 | 26.38 | 23/01/2018 |
|  | Cr2 | Cr1.2 | EUR | 41.69 | 26.38 | 23/01/2018 |
|  | Cr3 | Cr1.3 | EUR | 41.69 | 26.38 | 24/01/2018 |
|  | Cr4 | Cr1.4 | EUR | 41.69 | 26.38 | 23/01/2018 |
|  | Cr8 | Cr73.1 #4 | EUR | 37.89 | 22.73 | 22/01/2018 |
|  | Cr10 | Cr 79.9 #4 | EUR | 37.74 | 21.69 | 22/01/2018 |
|  | Cr11 | VIC-FR7-2 | EUR | 43.39 | 0.05 | 30/01/2018 |
|  | Cr12 | TOS-FR8-6 | EUR | 43.33 | 0.10 | 30/01/2018 |
|  | Cr14 | THE-GR2-5 | EUR | 40.64 | 22.94 | 23/01/2018 |
|  | Cr15 | NAB-PAL3-5 | ME | 32.25 | 35.22 | 29/01/2018 |

Accession ID = name given to accession in our study; Accession name =reference name of the accession in our database; Origin= geographical cluster: ASI = Asia, CASI = Central-Asia, EUR= Europe, ME= Middle-East.

**Table S2.** Comparison table of the number of fruits obtained in our studies (Present) and in past study from Petrone Mendoza *et al*. (Past).

| Species | Study | Competition | n | mean | sd | se | ci |
| --- | --- | --- | --- | --- | --- | --- | --- |
| Cbp | Present | alone | 144 | 1876.4306 | 1114.423 | 92.86859 | 183.57261 |
| Cbp | Present | compet | 144 | 1886.9514 | 1143.2137 | 95.26781 | 188.31514 |
| Cbp | Past | alone | 42 | 618.3571 | 374.745 | 57.82442 | 116.77878 |
| Cbp | Past | compet | 166 | 453.8193 | 366.5842 | 28.45245 | 56.17783 |
| Cg | Present | alone | 40 | 4630.075 | 2569.8053 | 406.32189 | 821.8636 |
| Cg | Present | compet | 40 | 4567.9 | 2758.1544 | 436.10251 | 882.10058 |
| Cg | Past | alone | 69 | 1516.971 | 1044.8369 | 125.78358 | 250.99723 |
| Cg | Past | compet | 273 | 941.2418 | 865.7293 | 52.39632 | 103.15388 |
| Co | Present | alone | 24 | 1745.9167 | 1257.287 | 256.64264 | 530.90575 |
| Co | Present | compet | 24 | 1424.375 | 865.6052 | 176.69092 | 365.51301 |
| Cr | Present | alone | 40 | 1745.975 | 1079.0003 | 170.60492 | 345.08103 |
| Cr | Present | compet | 40 | 1784.45 | 865.5654 | 136.85791 | 276.82125 |
| Cr | Past | alone | 60 | 915.75 | 542.0654 | 69.98034 | 140.03033 |
| Cr | Past | compet | 236 | 529.2161 | 381.2611 | 24.81799 | 48.89416 |

Species: Cbp, *Capselfia bursa-pastoris*; Cg, *C. grandiflora*; Co, *C. orientalis*; Cr, *C. rubella*. Data come from two different studies: the present study (Present) and a study carried out under controlled condition by Petrone Mendoza et al. in 2018 (Past). In both study the number of fruits was estimated in presence of competitors (comp.) or without competitor (alone). n= the number of individuals; mean = average number of flowers (/fruits); sd = standar deviation; se = standard error; ci= confidence interval

